# Recruitment of Archaeal DTD is a Key Event in the Emergence of Land Plants

**DOI:** 10.1101/2020.05.22.109876

**Authors:** Mohd Mazeed, Raghvendra Singh, Pradeep Kumar, Bakthisaran Raman, Shobha P. Kruparani, Rajan Sankaranarayanan

## Abstract

Land plant evolution is a major leap in the history of life that took place during the Neoproterozoic Era (∼800 Mya). Charophyceae, a class of rhyzophytic green algae emerged as a land plant with innovations in biochemical, cytological and developmental adaptations and played a crucial role in establishing life on the land^1,2^. One such striking architectural innovation is “root” that experience harsh environmental assaults such as floods, waterlogging and therefore is the epicentre for anaerobic fermentation, which produces toxic acetaldehyde^3^. Here, we show that such produced acetaldehyde makes N-ethyl-adducts on a central component of translation machinery aa-tRNA. The Plant kingdom is unique among life forms in possessing two chirality-based proofreading systems represented by D-aminoacyl-tRNA deacylases (DTD1 and DTD2), derived from Bacteria and Archaea^4^. We identified a unique role of archaeal derived chiral proofreading module DTD2 that selectively deacylates N-ethyl-D-aminoacyl-tRNAs (NEDATs) in plants. NEDAT deacylase function is exclusive to DTD2, as no other proofreading modules with similar substrates like canonical DTD1 and peptidyl-tRNA hydrolase (PTH) can clear NEDATs. Thus, the study elucidates the cause of hypersensitivity of DTD2 knockout plants for both ethanol and acetaldehyde. We further show NEDAT elimination is rooted in Archaea which possess the biosynthesis machinery for ethanol fermentation similar to plants. While absent in other algal branches, DTD2 can be identified in plants from land plant ancestors-Charophytes onwards. DTD2 is the only gene that has only archaeal origin among the genes ascribed for architectural and genomic innovations that happened in the land plant ancestors. The work has uncovered an important gene transfer event from methanogenic archaea to the charophytes in the oldest terrestrial ecosystem bog that contains excess of D-amino acids and deprived of oxygen.

---

Plants being sedentary in their lifestyle are dependent on roots for their nourishment and root has to survive in the oxygen-deprived environment of soil. The soil also being enriched with D-amino acids makes roots clutched with inescapable stress of D-amino acids compounded with anaerobic stress. The presence of D-amino acids in plants is inevitable and play vital roles in plant physiology such as the development of the pollen tube and also act as a major nitrogen source^5^. On the contrary, D-amino acids also cause cellular toxicity by generating D-aminoacyl-tRNAs (D-aa-tRNAs) which arrests free tRNA pool from translation. Removal of mischarged D-amino acids from D-aa-tRNAs is termed as chiral proofreading^6^ that enables the perpetuation of homochirality in the cellular proteome^7^. D-amino acid editing modules are universally conserved; D-aminoacyl-tRNA deacylase 1 (DTD1) is present in both Bacteria and Eukaryotes and DTD2 is in Archaea and plants^4^. Plants are the only organisms that possess both the evolutionarily distinct chiral proofreading modules DTD1 and DTD2^8,9^. Surprisingly, DTD2 knock out plants show enhanced sensitivity to both ethanol and acetaldehyde, which are metabolites of ethanol fermentation^10,11^. It has been a long-standing and intriguing question that how D-amino acid detoxifying system-DTD2 confers protection to plants from toxic intermediates of incessant ethanol fermentation especially acetaldehyde during anaerobic stress.

## Acetaldehyde creates ethyl modification on aa-tRNA

DTD2 is known to act on D-aminoacyl-tRNAs and therefore we set out to investigate whether DTD2’s ability to protect plants from acetaldehyde is mediated through aa-tRNAs. We treated aa-tRNAs with acetaldehyde followed by a reducing agent (Fig: 1A). Surprisingly, this resulted in the shift of the position of aa-tRNA corresponding spot on the thin-layered chromatography (TLC) (Fig: 1B). The ester linkage between amino acid and tRNA of acetaldehyde modified aa-tRNA (half-life: 13 minutes) showed enhanced stability compared to aa-tRNA (half-life: 1 minute) up on alkali-treatment (Fig: 1C). It has been shown that N-acetyl-L-aa-tRNAs and peptidyl-tRNAs are more stable compared to aa-tRNAs^12^, and they are hydrolysed by peptidyl-tRNA hydrolase (PTH) (Fig: 2F, G). Our assays revealed that acetaldehyde treated aa-tRNAs are much more stable than acetyl-aa-tRNAs (half-life: 5 minutes) (Fig: 1C). Ultra-stability makes N-blocked aa-tRNAs more lethal to the cells because they permanently sequester free tRNA pool, which is evident in the case of PTH knockout of *E. coli*^13,14^. These observations indicated that acetaldehyde can modify aa-tRNA. To characterize the chemical nature of the modification, electrospray ionization mass spectrometry (ESI-MS/MS) of both acetaldehyde treated and untreated non-hydrolysable analogs of aminoacyl-tRNA was performed. These analogs represent both amino acid and tRNA part of aa-tRNA, where the amino acid is bound to terminal adenosine-76 of tRNA with non-hydrolysable amide bond. Mass spectrometry data clearly showed that modification created by acetaldehyde on aa-tRNA is ‘ethyl’ moiety, which is a covalent modification (Fig: 1D). Incubation of acetaldehyde with aminoacyl-tRNAs and analogs bearing amino acids with different chirality (L- and D-) and varied side chains (Phe-tRNA^Phe^ Tyr-tRNA^Tyr^, Ala-tRNA^Ala^, Ser-tRNA^Ser^ and Thr-tRNA^Thr^) (Tyr2AA, Ala3AA, Val3AA and Thr2AA) revealed that acetaldehyde modification is independent of chirality and side-chain chemistry of aa-tRNAs (Fig: 1B, E) (Extended Data Fig: 1, 3). Since acetaldehyde is known to target free amino groups, to know the site of ethyl modification on the analog, tandem mass spectrometry fragmentation (MS^2^) of ethyl-modified analogs was performed (Fig: 1G, H) (Extended Data Fig: 2, 3). Fragmentation experiments (MS^2^) confirmed that modifications happen only on the amino acid part of aa-tRNA (Fig: 1F, G, H) (Extended Data Fig: 2, 3) but not on tRNA even at 1000-fold molar excess of acetaldehyde (Extended Data Fig: 6D, H) (Extended Data Fig: 7A, D) (Extended Data Fig: 8A, D). These experiments clearly established the unknown link between anaerobic stress metabolite acetaldehyde and a key constituent of the translation machinery aa-tRNA. It also prompted us to hypothesise that, DTD2 relieves anaerobic stress by deacylating ethyl modified amino acids from N-ethyl-aa-tRNAs (NEATs) in plants.

**Fig 1:**
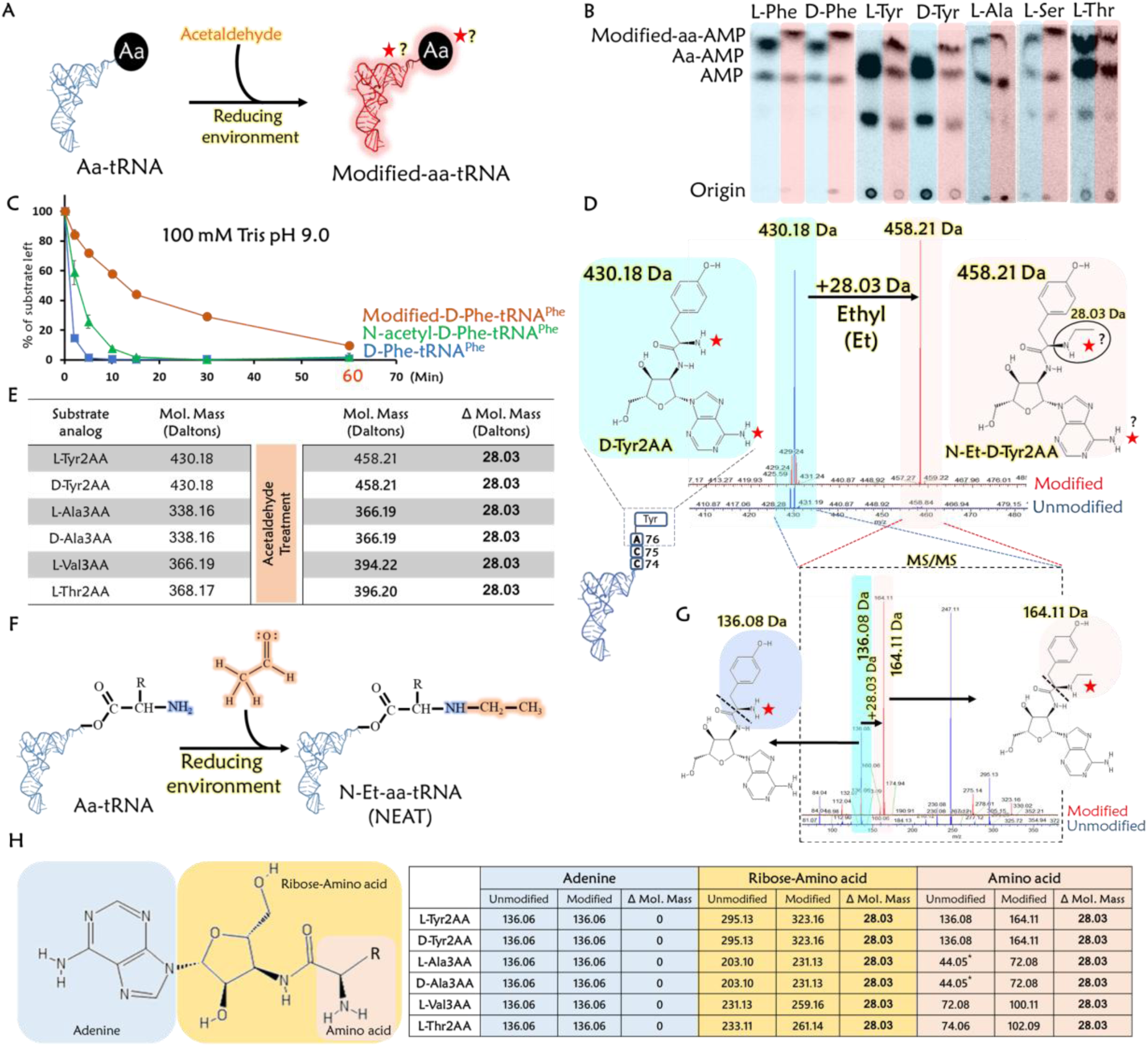
Acetaldehyde targets amino acid of aa-tRNA and generates ethyl modification. (A) Schematic showing reaction of aa-tRNA and acetaldehyde in the presence of a reducing agent. (B) TLC showing differences in the spot positions of aa-AMP corresponding to unmodified (blue) and acetaldehyde modified (pink) L-Phe-tRNA^Phe^, D-Phe-tRNA^Phe^, L-Tyr-tRNA^Tyr^, D-Tyr-tRNA^Tyr^, L-Ala-tRNA^Ala^, L-Ser-tRNA^Ser^, and L-Thr-tRNA^Thr^. (C) Alkali treatment of D-Phe-tRNA^Phe^, acetaldehyde modified-D-Phe-tRNA^Phe^ and N-acetyl-D-Phe-tRNA^Phe^ showing ultra-stability of acetaldehyde modified-D-Phe-tRNA^Phe^. (D) ESI-MS analysis showing ethyl modification (a clear shift of 28.03 Da) on acetaldehyde treated substrate analog of D-Tyr-tRNA^Tyr^ (D-Tyr2AA) (unmodified DY2AA with m/z=430.18 (blue) and modified DY2AA with m/z=458.21 (pink)). (E) Table showing m/z peaks observed in ESI-MS analysis of acetaldehyde treated analogs of multiple aa-tRNAs such as L-Tyr2AA (L-Tyr-tRNA^Tyr^), D-Tyr2AA (D-Tyr-tRNA^Tyr^), L-Ala3AA (L-Ala-tRNA), D-Ala3AA (D-Ala-tRNA), L-Val3AA (L-Val-tRNA) and L-Thr2AA (L-Thr-tRNA) confirmed variations in the side chain and chirality did not influence the modification. (F) Schematic shows acetaldehyde creates ethyl modification on aa-tRNA. (G) Fragmentation experiments (MS^2^) showing adenosine and tyrosine peaks in unmodified (blue) DY2AA, adenine and ethyl modified (pink) tyrosine corresponding peaks in modified DY2AA. (H) Table showing m/z peaks observed during MS^2^ analysis, adenine and amino acid (immonium ion) in unmodified analogs (blue), adenine and ethyl modified amino acid alone and along with ribose in modified (pink) analogs of L-Tyr-tRNA^Tyr^ (L-Tyr2AA), D-Tyr-tRNA^Tyr^ (D-Tyr2AA), L-Ala-tRNA (L-Ala3AA), D-Ala-tRNA (D-Ala3AA), L-Val-tRNA (L-Val3AA) and L-Thr-tRNA (L-Thr2AA) (* indicates expected molecular weights) (related mass spectrometry data is summarised in Extended Data Fig: 3).

**Fig 2:**
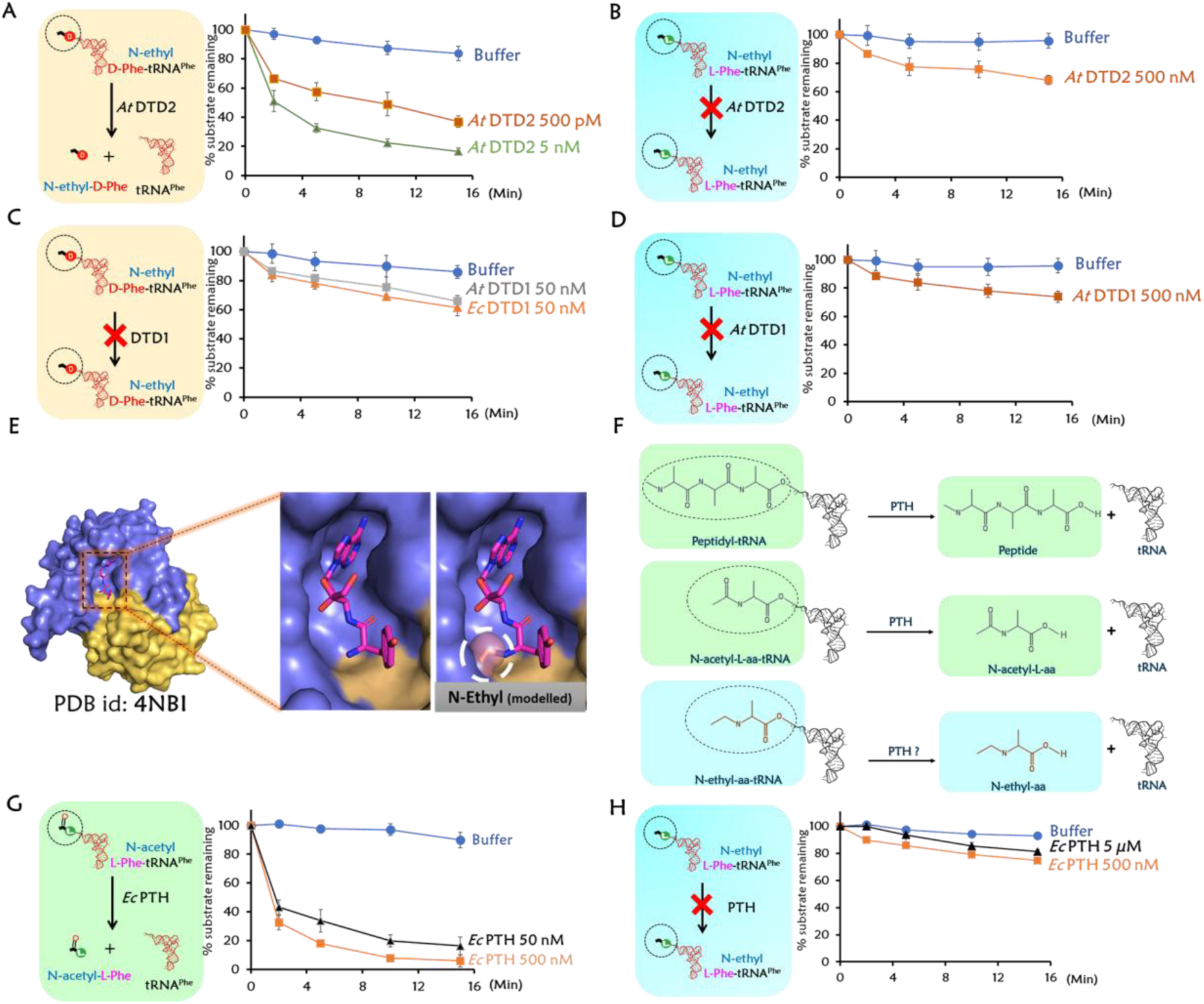
*At* DTD2 readily acts on NEDATs but not DTD1 and PTH. (A) N-ethyl-D-Phe-tRNA^Phe^ (*At*) and (B) N-ethyl-L-Phe-tRNA^Phe^ (*At*) deacylations with *At* DTD2. (C) Both prokaryotic and eukaryotic DTD1 did not deacylate N-ethyl-D-Phe-tRNA^Phe^. (D) Deacylation of N-ethyl-L-Phe-tRNA^Phe^ with *At* DTD1. (E) Crystal structure of DTD1 in complex with D-Tyr3AA and ethyl modification on the α-NH_2_ group of D-tyrosine of the ligand (Model). (F) Peptidyl-tRNA, N-acetyl-aa-tRNA and N-ethyl-aa-tRNA are chemically very subtle. *Ec* PTH deacylated N-acetyl-L-Phe-tRNA^Phe^ (*Ec*) (G) but not N-ethyl-L-Phe-tRNA^Phe^ (*Ec*) (H).

## DTD2 alleviates anaerobic stress in plants by recycling NEDATs

To check this possibility, *in vitro* deacylation assays were performed by incubating DTD2 of *Arabidopsis thaliana* (*At* DTD2) with both N-ethyl-L-aa-tRNAs (NELATs: N-ethyl-L-Phe-tRNA^Phe^ (*At*)) and N-ethyl-D-aa-tRNAs (NEDATs: N-ethyl-D-Phe-tRNA^Phe^ (*At*)). Interestingly, *At* DTD2 readily deacylated NEDATs to N-ethyl-D-amino acids and free tRNAs but not NELATs (Fig: 2A, B) (Extended Data Fig: 4). The activity of DTD2 on NEDATs is referred to as ‘NEDAT deacylase (ND) activity’ in the rest of the article. Since plants also possess canonical chiral proofreading module DTD1, we performed deacylations of NELATs and NEDATs with DTD1. Both the Bacterial (*Ec*) and Eukaryotic (*At*) DTD1s did not act on NELATs and also on NEDATs (Fig: 2C, 2D) (Extended Data Fig: 5). DTD1’s lack of activity on NEDATs is further corroborated by the earlier crystal structure of DTD1 in complex with substrate analog D-Tyr3AA^6,15^. Wherein the D-amino acid of D-aminoacyl-tRNA snugly fits into the active site of DTD1, thereby sterically excludes N-ethylated amino group of D-amino acid (Fig: 2E). PTH, as mentioned above acts on N-acetyl-L-aa-tRNAs and these acetylated substrates are similar in size but chemically distinct from ethylated substrates (Fig: 2F). Therefore, we expected that PTH could act on NELATs, but, to our surprise, PTH had no activity on NELATs (Fig: 2G, H) (Extended Data Fig: 4E) suggesting the strict chemical selectivity of PTH. These findings on the biochemical activity of DTD2 led to the identification of its distinct role as a unique proofreader of D-aa-tRNA adducts. As mentioned above, acetaldehyde also modified L-aa-tRNAs as efficiently as D-aa-tRNAs but none of the enzymes viz, DTD1, DTD2 and PTH could deacylate the NELATs (Fig: 1B, E) (Extended Data Fig: 1, 3) (Fig: 2B, D, H). Elongation factor thermo unstable (EF-Tu) binds to D-aa-tRNAs with 25-fold lesser affinity compared to L-aa-tRNAs. While nearly 90% of the L-aa-tRNAs are EF-Tu bound and are used at the ribosome by participating in protein synthesis, the D-aa-tRNAs are rejected by the cellular chiral checkpoints^16,17^. Therefore, particularly D-aa-tRNAs are susceptible to modification by the freely diffusing acetaldehyde. In addition, DTD2’s ability to rescue acetaldehyde toxicity in *Arabidopsis* in combination with its lack of activity on NELATs highlight that NEDATs are the cellular substrates of DTD2. Overall, the above findings clearly show that DTD2 alleviates the effect of acetaldehyde by actively clearing NEDATs during anaerobic stress.

## ND activity is rooted in the Archaea

Apart from plants, DTD2 is also conserved in Archaea (Fig: 3A)^8,9^ and therefore we further checked whether archaeal DTD2 possess the ND activity. We performed adduct deacylation experiments with distantly related archaeal DTD2s from the organisms *Pyrococcus horikoshii (Pho* DTD2*), Methanocaldococcus jannaschii (Mj* DTD2) and *Archaeoglobus fulgidus (Af* DTD2). Remarkably, all three archaeal DTD2s acted on NEDATs with the same efficiency as plant DTD2s (Fig: 3C, D, E). Similar to plant DTD2s, archaeal DTD2s also possess the chiral selectivity towards adducts by not acting on NELATs (Fig: 3F) (Extended Data Fig: 4). The presence of DTD2 across archaea and the conservation of ND activity highlight that DTD2’s ability to clear NEDATs is ‘rooted’ in the archaeal branch of life. Therefore it is clear that DTD2 is acquired from Archaea during the evolution of plants and completely absent in other major eukaryotic forms of life i.e. opisthokonts, that include fungi and animals.

**Fig 3:**
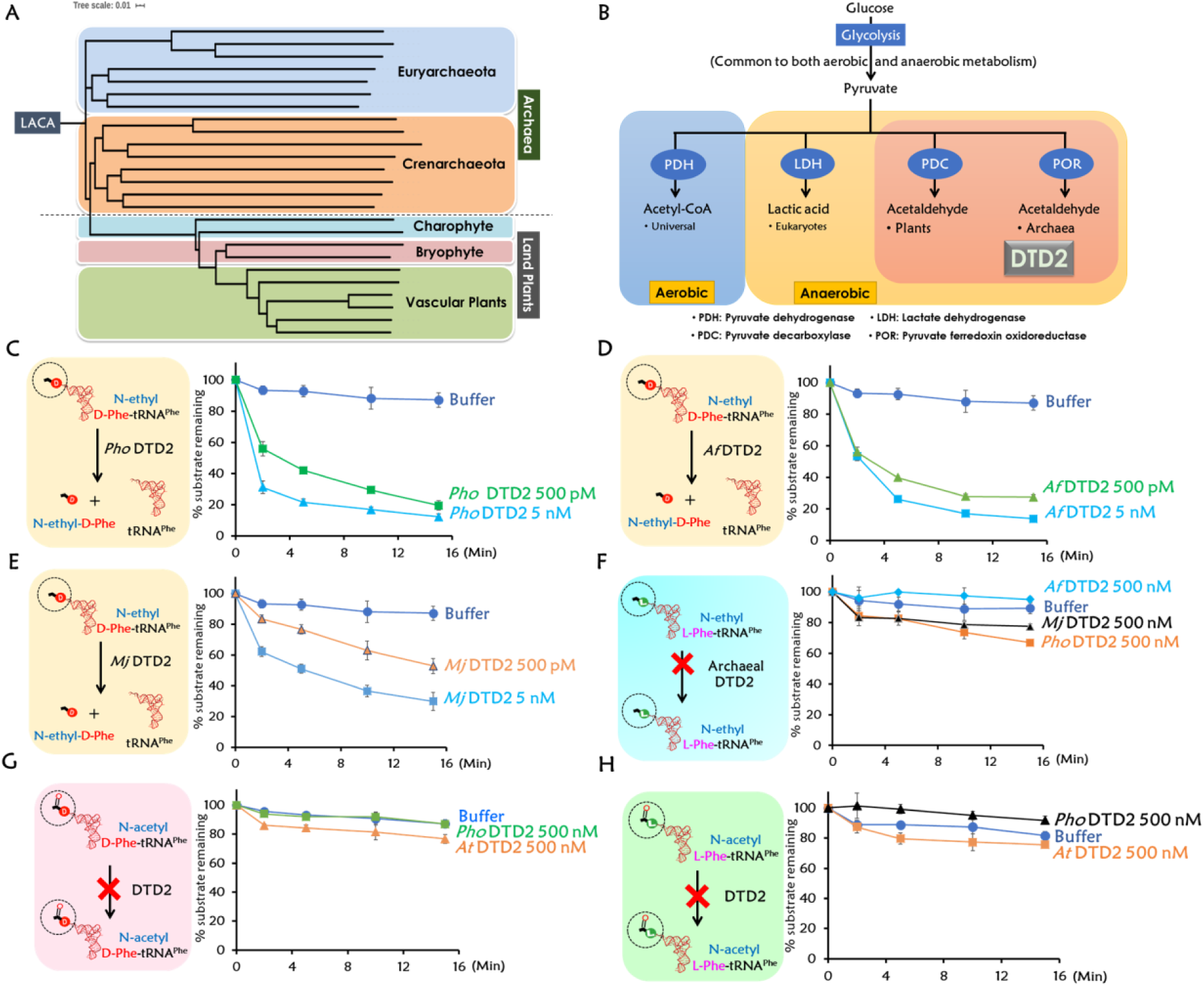
DTD2’s ND activity and chiral selectivity are conserved throughout Archaea. (A) Phylogenetic classification of archaeal and plant DTD2. (B) Co-occurrence of DTD2 with acetaldehyde biosynthesis i.e., only in Archaea and plants. Deacylations of N-ethyl-D-Phe-tRNA^Phe^ (*Pho*) with *Pho* DTD2 (C), *Af* DTD2 (D), and *Mj* DTD2 (E). (F) N-ethyl-L-Phe-tRNA^Phe^ (*Pho*) is not deacylated with *Pho, Af* and *Mj* DTD2 even at 500 nM enzyme concentration. Deacylations of N-acetyl-D-Phe-tRNA^Phe^ (*Pho*) (G) and N-acetyl-L-Phe-tRNA^Phe^ (*Pho*) (H) with both archaeal (*Pho)* and plant (*At)* DTD2.

We next interrogated substrate specificity of DTD2 in terms of modification by performing deacylations with N-acetyl-D-aa-tRNAs and found that both, archaeal (*Pho*) and plant (*At*) DTD2s did not hydrolyse acetyl substrates (Fig: 3G, H) suggesting, DTD2 is specific for ethyl modification and evolved exclusively for ND activity. It is therefore intriguing to probe what physiological constraints necessitated the conservation of DTD2 and its ND activity in Archaea and plants. To this end, we searched for common physiological and biochemical characteristics among Archaea and plants. Strikingly, we noted that acetaldehyde biosynthesis machinery is indeed conserved exclusively in Archaea and plants^3,18,19^. In Archaea, pyruvate ferredoxin oxidoreductase (POR) converts pyruvate to acetaldehyde and pyruvate decarboxylase (PDC) performs the same function in plants (Fig: 3B). These machineries are mostly absent in Bacteria and Opisthokonta^20,21^. This analysis provides a strong biochemical basis for the requirement of DTD2 in Archaea and plants.

## DTD2 distribution is exclusive to land plants and their ancestors in the Plant kingdom

We then set out to identify when plants acquired the archaeal version of DTD i.e. DTD2. Extensive bioinformatic analysis of DTD2 distribution in plants revealed that DTD2 is strictly confined to land plants and their ancestors Charophytes. While DTD2 is completely absent in different clades of Algae, which include Glaucophytes, Red algae and Green algae, however, recently sequenced genomes of Charophytes-*Klebsormidium nitens*^22^ and *Chara braunii*^23^ encode DTD2 (Fig: 4A). Structure-based sequence alignment of DTD2s including *K. nitens* and *C. braunii* revealed that both the Charophyte DTD2s conserve the residues important for catalysis (Fig: 4B)^9^. Recent reports identified Charophytes as land plant ancestors that acquired many land plant-specific characteristics^1,2,22,23^. Genes of phytohormone biosynthesis and their receptors, multicellularity, Skp, Cullin, F-box containing (SCF) complex and cyclic electron flow (CEF) are land plant-specific adaptations appeared just in Charophytes, conserved across land plants and completely absent in other clades of the Algae^22^. Similar presence of DTD2 from Charophytes to land plants clearly suggests that its acquirement is an important land plant-specific adaptation. We searched for the origins of the above-mentioned genes that are responsible for the adaptation of land plants. Except DTD2, these genes have either only Bacterial origin, both Bacterial and Archaeal origin or appeared *de novo* as genomic innovations in Charophytes (Fig: 4C) (Extended Data Fig: 11). DTD2 is reported to be one of the two proteins unique to Archaea and plants; the other protein is topoisomerase VI subunit B (TopVIB)^11^. Mutations in TopVIB affect cell proliferation and endoreduplication, which are unique characteristics of plants ^24–27^. While TopVIB is a subunit of archaeal type VI topoisomerase, DTD2 is a complete functional protein. Unlike DTD2, TopVIB is conserved in all the lineages of Algae apart from the land plants (Extended Data Fig: 10). Overall, this suggests DTD2 is the only protein unique to Archaea and land plants. A thorough phylogenetic analysis of all the known archaeal and land plant DTD2 highlighted that DTD2 of *K. nitens* and *C. braunii* are closer to the archaeal ones than any other land plant DTD2 (Fig: 3A). Among archaeal DTD2, methanogenic archaea (*Methanocella conradii*) shares maximum identity with most of the DTD2 of the land plants (Fig: 4D) (Extended Data Fig: 10); in agreement, *Methanocella* is found to be in symbiotic association with lower plants in bogs. Bogs are the oldest terrestrial ecosystem deprived of oxygen and enriched with D-amino acids^28^. This emphasises on a single acquisition event of DTD2 from methanogenic archaea to the Plant kingdom that happened at the basal radiation of the land plants Charophytes during Neoproterozoic Era (∼800 Mya). As mentioned earlier, except Charophytes, DTD2 is absent in all the lineages of Algae, which further strengthens the above argument that DTD2 is acquired from Archaea by the Charophyta, which facilitated plants to colonise and thrive on the land. One of the important architectural innovations that happened during the evolution of land plants is the ‘root’, which is the primary organ exposed to higher anaerobic stress and excess D-amino acids in the soil^3,5,29–31^. This is well correlated with higher expression of DTD2 in the roots compared to the other organs^11^. The confluence of these two independent stresses intensifies the formation of NEDATs and necessitates the acquirement of DTD2 from Archaea.

**Fig 4:**
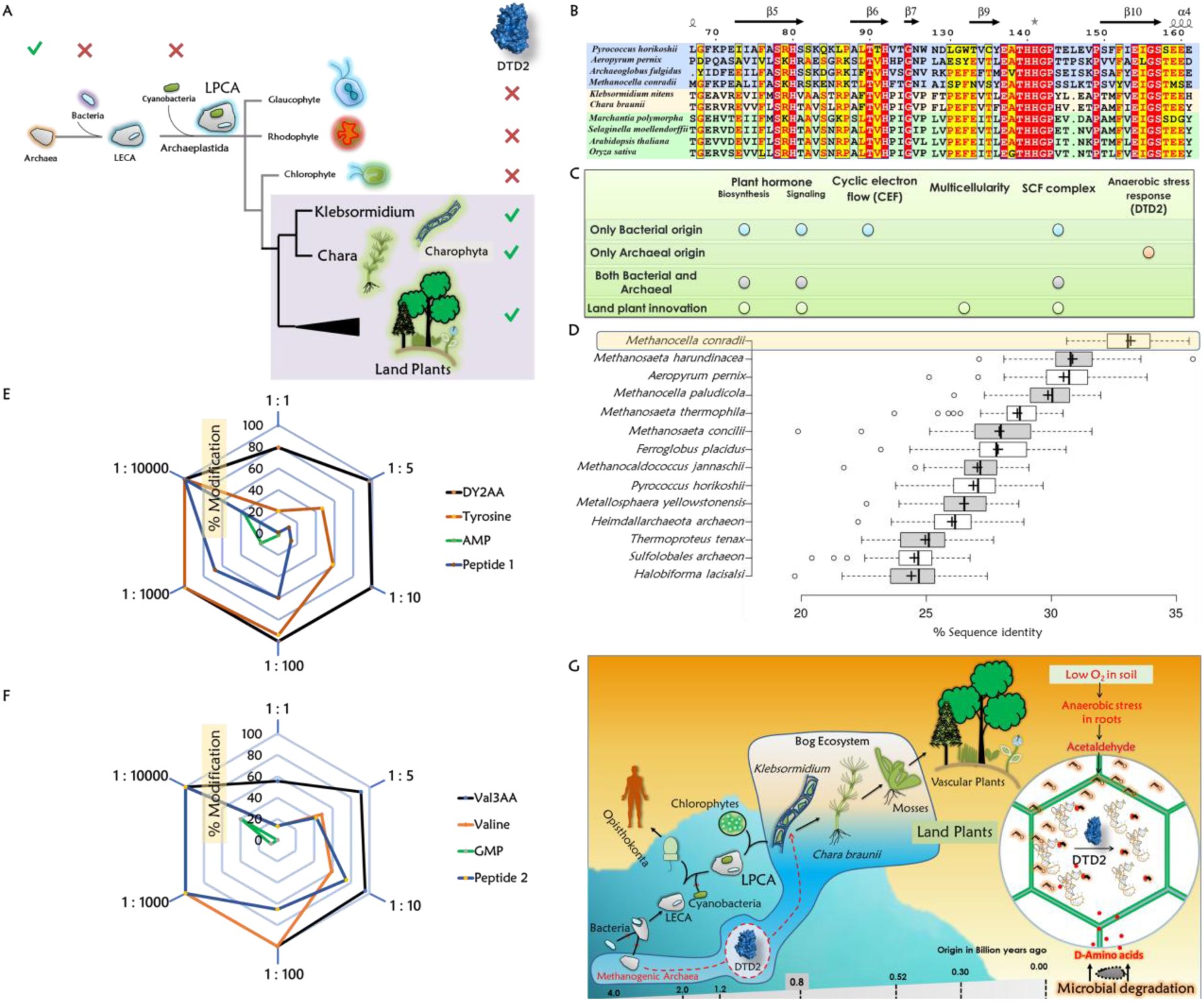
Acquiring archaeal DTD2 is a key event in the evolution of land plants to mitigate the complexity created by the confluence of anaerobic stress and D-amino acids. (A) DTD2 distribution is seen only in Archaea, land plants and their ancestors; DTD2 is completely absent in multiple lineages of Algae. (B) Structure-based sequence alignment shows conservation of active site residues in *K. nitens* and *C. braunii*. (C) DTD2 is the gene of only archaeal origin that is responsible for land plant-specific innovation. (D) Boxplot displays DTD2 of *M. conradii* shares maximum identity (33.18%) with DTD2 of most of the plants, the horizontal axis represents sequence identities with all 80 plant DTD2s and the vertical axis shows the representative archaeal sequences compared with all plants (Complete list is provided supplementary table S3). (E, F) Radar charts showing aa-tRNAs are hypersensitive to acetaldehyde compared to amino acids, peptides and nucleotides (Each corner represents the ratio of acetaldehyde used and hexagonal rings depict extant of modification). (G) Model showing DTD2 recruitment in Charophytes and its implications in land plant evolution.

## Aa-tRNA is the first target to get modified by acetaldehyde

Chronic alcohol consumption in humans causes various cancers^32-34^. The causative agent for the devastating effects of alcohol consumption is found to be acetaldehyde and thus it is demonstrated as a genotoxic and teratogenic agent in mice and humans^34^. As observed with aa-tRNA adducts, acetaldehyde also makes a nucleophilic attack on free amino groups of DNA and proteins that creates permanent ethyl adducts^35^. To compare relative acetaldehyde sensitivities of free amino acids, proteins and nucleotides with that of aa-tRNA, when treated with acetaldehyde, we found that aa-tRNAs are more prone to modification than other counterparts. Since 10000-fold molar excess of acetaldehyde is required to modify nucleotides, suggests that DNA is relatively very less prone to modification. Amino acids and peptides are found to be second preferable targets for acetaldehyde. Interestingly, equimolar acetaldehyde is enough to modify aa-tRNAs (Fig: 4E, F) (Extended Data Fig: 6-9). This clearly suggests that during anaerobic stress or acetaldehyde burst aa-tRNA is the first target that undergoes modification. Toxic effects of acetaldehyde are quenched by multiple tiers of detoxifying systems especially acetaldehyde dehydrogenase (ALDH) and Fanconi anaemia group D2 protein (FANCD2). ALDH and FANCD2 do not act directly on ethyl modified substrates^32–34^, whereas DTD2 is the only enzyme that directly removes N-ethyl-amino acids from NEDATs. Since aa-tRNAs are the primary targets for acetaldehyde, DTD2 is expected to provide an early response during anaerobic stress.

Overall, our study demystifies the link between DTD2 and its role in protecting plants from acetaldehyde toxicity. The work provides the first empirical evidence that acetaldehyde targets aa-tRNAs and also identifies a unique role of DTD2 as a proofreader of D-aa-tRNA adducts. The strong correlation between DTD2 and acetaldehyde biosynthesis along with the presence of DTD2 only in land plants and their ancestors clearly suggest that acquiring archaeal DTD2 is a key event in the emergence and evolution of land plants (Fig: 4G).

## Acknowledgments

The authors acknowledge Dr. R. Nagaraj, CSIR-CCMB for fruitful discussions and providing peptides. Dr. S. Raghavan, CSIR-IICT for insightful discussions. M. M. thanks Department of Biotechnology, India for research fellowship. R. S. thanks SERB-NPDF. P. K. thanks CSIR for research fellowship. R. S. N. acknowledges healthcare theme project, CSIR, India, J.C. Bose Fellowship of SERB, India, and Centre of Excellence Project of Department of Biotechnology, India.

## Author contributions

M.M., R.S., P.K., S.P.K. and B.R. designed and performed the experiments. R.S.N. conceived and supervised the study. All the authors analyzed the data. M.M., R.S. and R.S.N. wrote the manuscript with the help from P.K. and all the authors reviewed it.

## Materials and Methods

### Materials

Materials were obtained from Merck unless otherwise mentioned. ESI-MS was performed by using Thermo Scientific Q-Exactive mass spectrometer. Plasmid coding for different subunits of phenylalanyl-tRNA synthetase (PheRS), pKPY514 was a gift from David Tirrell (Addgene plasmid #62598; http://n2t.net/addgene:62598 ; RRID:Addgene_62598) ^36^. Superdex 75 and Sulfopropyl-sepharose columns were purchased from GE Healthcare Life Sciences, USA. The non-hydrolysable analogs of aa-tRNAs were purchased from Jena Biosciences, Germany.

### Generation of adducts on aa-tRNAs and its analogues

Adducts on aa-tRNAs and their non-hydrolysable analogs were generated by incubating 2 µM aa-tRNA and 100 µM aa-tRNA analog with 20 mM acetaldehyde at 37°C for 30 minutes. To remove excess acetaldehyde from the reaction, tubes were subjected to Eppendorf 5305 Vacufuge plus Concentrator. Above mixture was reduced with 400 mM NaCNBH_3_ and incubated at 37°C for 30 minutes. All the reactions were performed on a shaking incubator with 300 RPM. Acetaldehyde treated analogs were characterized using mass spectrometry without any further processing, whereas acetaldehyde modified aa-tRNAs were ethanol precipitated for overnight to remove salts and metal ions. Pellets were re-suspended with 5 mM sodium acetate (pH 5.4). Acetaldehyde modification of aa-tRNA was assessed and quantified by spotting on cellulose F (Merck) TLC plates^6^.

### Probing relative acetaldehyde sensitivities of biomolecules

200 µM of each of non-hydrolysable anlogs (DY2AA & Val3AA), amino acids (D-Tyr & L-Val), peptides (Pep 1 & Pep 2) and nucleotides (AMP & GMP) were incubated with different concentrations of acetaldehyde (200 µM, 1 mM, 2 mM, 20 mM, 200 mM, and 2 M) along with 400 mM NaCNBH_3_ and incubated at 37°C for 30 minutes. The tubes were subjected to Eppendorf 5305 Vacufuge plus Concentrator. All the samples were characterized using mass spectrometry without any further processing.

### Cloning, expression and purification

Genes encoding DTD2s of *Pyrococcus horikoshii* (*Pho*), *Aeropyrum pernix* (*Ap*), *Methanococcus jannaschii* (*Mj*), *Archaeoglobus fulgidus* (*Af*) and tyrosyl-tRNA synthetase (TyrRS) of *Thermus thermophilus* were PCR amplified from their genomic DNA using forward and reverse primers. *Arabidopsis thaliana* (*At*) DTD2 gene was amplified from the cDNA of *A. thaliana*. Genes were cloned into the pET28b vector by using restriction-free cloning^37^. Primers used in this study are listed in the table 1. *Pho* DTD2, *Af* DTD2, *At* DTD2, *T*t*h* TyrRS were transformed and overexpressed into *E. coli* BL21(DE3), *Ec* PheRS into *E. coli* M15, **and** *Mj* DTD2 into BL21CodonPlus (DE3)-RIL™strain of *E. coli*. Purification of 6X His-tagged proteins (*Mj* DTD2-C-His, *At* DTD2-N-His, *T*t*h* TyrRS-N-His and *Ec* PheRS-N-His) was performed by Ni-NTA affinity chromatography, followed by size exclusion chromatography (SEC). SEC was performed by using a Superdex 75 column (GE Healthcare Life Sciences, USA). Untagged proteins (*Pho* DTD2-No-tag and *Af* DTD2-No-tag) were purified using cation exchange chromatography (CEC) followed by SEC. For CEC, cultures were lysed in buffer containing 50 mM Bis-Tris (pH 6.5) and 20 mM NaCl, lysed cultures were heated at 70°C for 30 minutes before subjecting to centrifugation (18000 RPM for 30 minutes at 4°C). The supernatant was subjected to CEC column and proteins were eluted with a gradient of NaCl from 50 mM to 200 mM. Sulfopropyl-sepharose (GE Healthcare Life Sciences, USA) column was used for CEC. Buffer containing 100 mM Tris (pH 8.0), 200 mM NaCl, 5 mM 2-mercaptoethanol (β-ME), and 50% Glycerol was used to store all the purified DTD2 proteins.

### Sample preparation and mass spectrometry (MS)

Samples were dissolved in water containing 10% methanol and 1% acetic acid and were subjected to ESI-mass spectrometry using a Q-Exactive mass spectrometer (Thermo scientific) by infusing through heated electrospray ionization (HESI) source operating at a positive voltage of 3.5 kV. The method of targeted selected ion monitoring (t-SIM) (with an isolation width of 2 m/z and the inclusion lists of the theoretical m/z of the MH+ ion species) was used to acquire the mass spectra (at the resolving power of 70000@200m/z). The high energy collision-induced (HCD) MS-MS spectra with the normalized collision energy of 25 of the selected precursor ion species specified in the inclusion list (having the actually observed m/z value from the earlier t-SIM analysis) were acquired using the method of t-SIM-ddMS2 (at the isolation window of 1 m/z or 0.5 m/z (in cases of AMP and GMP) at the ddMS2 resolving power of 35000@200m/z).

### *In vitro* biochemical experiments

*E. coli* tRNA^Phe^, *P. horikoshii* tRNA^Phe^, and *A. thaliana* tRNA^Phe^ were generated by using the MEGAshortscript T7 Transcription Kit (Thermo Fisher Scientific, USA). tRNAs were end-labelled with [α-^32^P] ATP (BRIT-Jonaki, India) using CCA-adding enzyme^38^. Phenylalanylation of tRNA^Phe^ was done by incubating 1 µM of tRNA^Phe^ with 2 µM *E. coli* PheRS in a buffer containing 100 mM HEPES (pH 7.5), 10 mM KCl, 30 mM MgCl2, 50 µM D-Phe and L-Phe, and 2 mM ATP at 37°C for 15 min. Deacylation experiments were carried as mentioned earlier ^6^. All the experiments were performed in triplicates. Mean values were used to plot the graphs and error bars denote the standard deviation from the mean value of three independent observations.

### Bioinformatic analysis

DTD2 protein sequences were retrieved from NCBI (https://www.ncbi.nlm.nih.gov/) and were subjected to phylogenetic tree construction. The online version of Clustal Omega (https://www.ebi.ac.uk/Tools/msa/clustalo/)and iTOL web server (http://itol.embl.de) were used to construct a phylogenetic tree. The list of plants and archaea, whose genomes completely sequenced were obtained from KEGG GENOME database (http://www.genome.jp/kegg/genome.html) and in addition, *Klebsormidium nitens* and *Chara braunii* were also included in this study. Percentage identity table was generated for 80 plant and 140 archaeal sequences by using Clustal Omega. Box plot was plotted with percentage identity to all plant DTD2 sequences on the horizontal axis and respective archaea on the vertical axis. For control, 290 plant and 500 archaeal sequences of Topo6B were retrieved from UniProt blast using *Arabidopsis* Topo6B as a query. All Topo6B sequences were used for box plot analysis.T-coffee (http://tcoffee.crg.cat/) server was used to prepare structure-based multiple sequence alignment of DTD2 and the corresponding figure was generated using ESPript 3.0 (http://espript.ibcp.fr/ESPript/cgi-bin/ESPript.cgi). To figure out the origins of genes/processes, which are essential for terrestrialization of plants, we retrieved the protein sequences of the genes involved in various processes (originated in *Klebsrmidium*) from NCBI (https://www.ncbi.nlm.nih.gov/) and manually performed protein BLAST-based in silico search against genome sequences of Bacteria and Archaea. We used query coverage (70%) and percentage identity (20) as a cutoff.

## Extended Data Figures

**Extended Data Fig 1:**
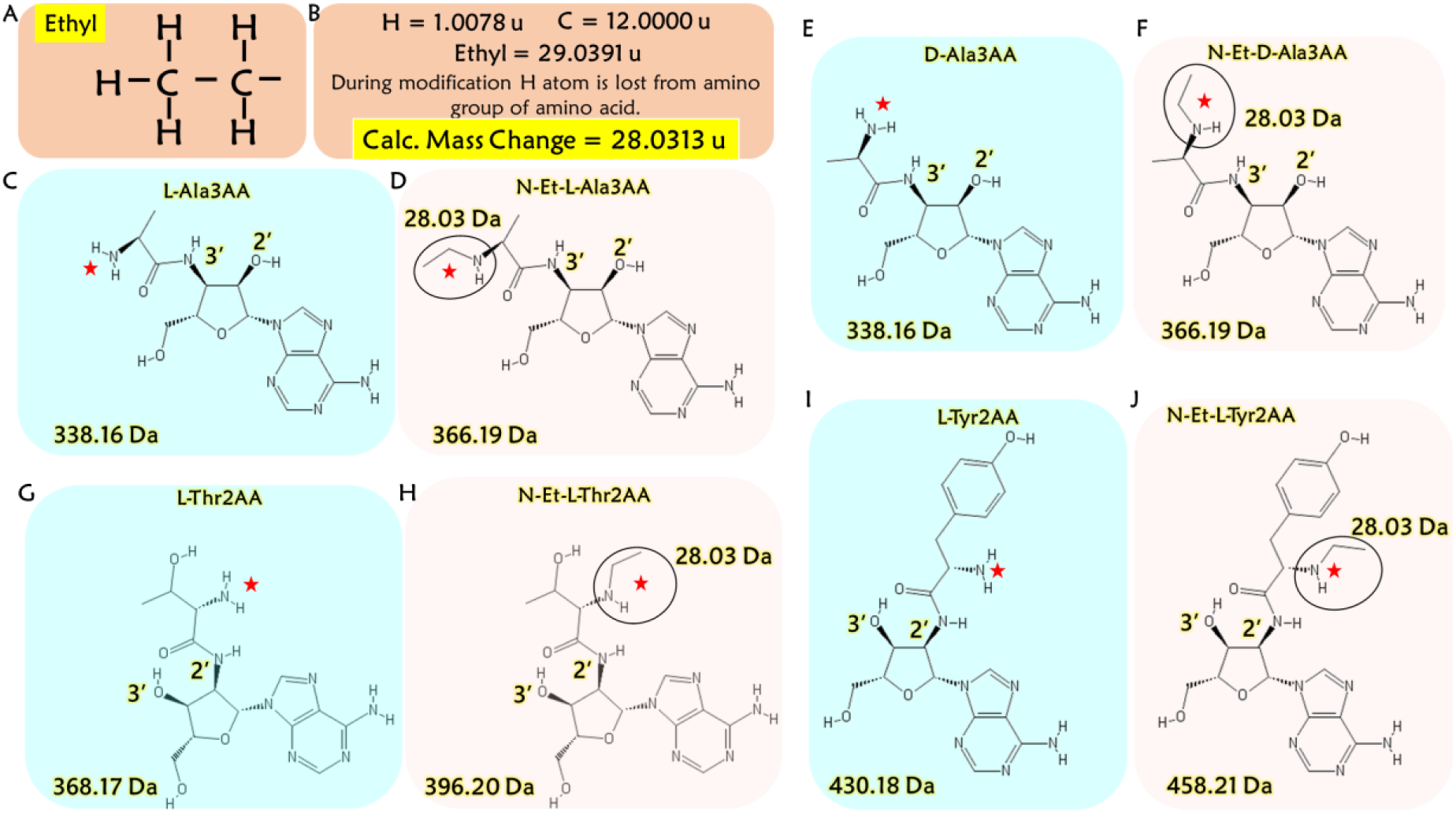
Structures and calculated molecular weights of acetaldehyde treated and untreated analogs. (A) Chemical structure of ethyl moiety. (B) The calculated molecular weight of ethyl modification. (C) Unmodified L-Ala3AA (D) modified L-Ala3AA (E) unmodified D-Ala3AA (F) modified D-Ala3AA (G) unmodified L-Thr2AA (H) modified L-Thr2AA (E) unmodified L-Tyr2AA (D) modified L-Tyr2AA.

**Extended Data Fig 2:**
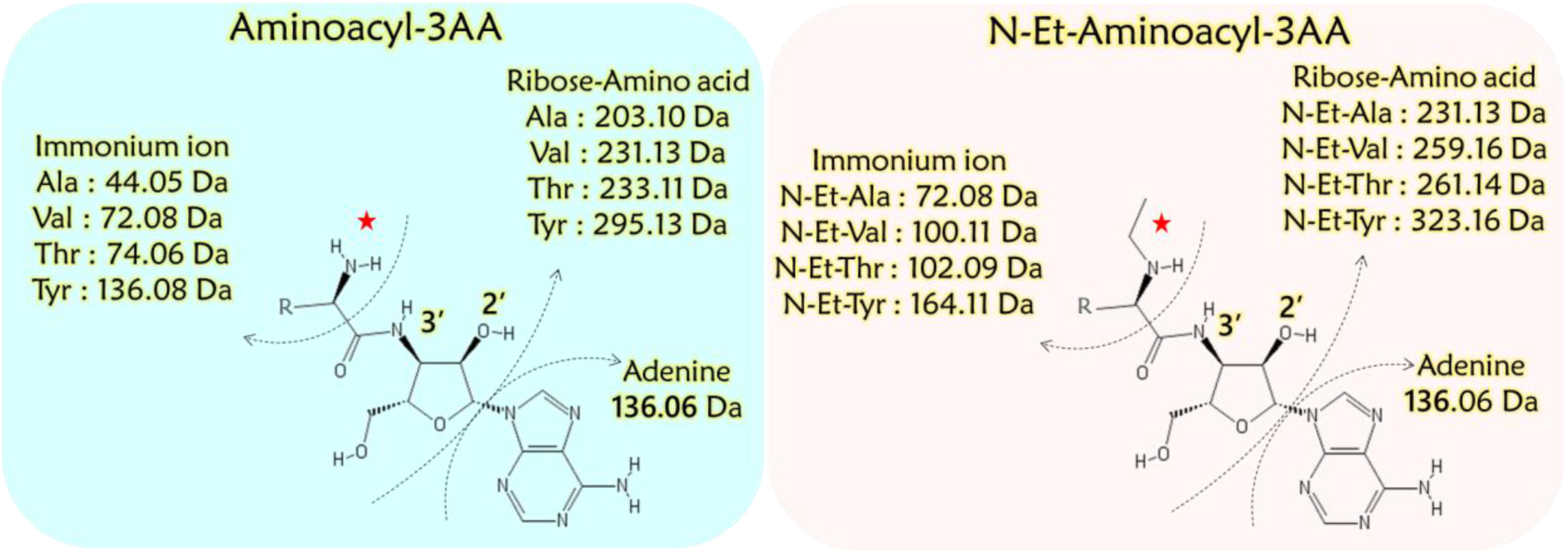
Tandem mass spectrometry fragmentation profile (MS^2^) of acetaldehyde treated and untreated analogs. Expected fragment size and mass for labelled peaks observed during fragmentation of various aminoacyl-tRNA analogs shown in Fig S3.

**Extended Data Fig 3:**
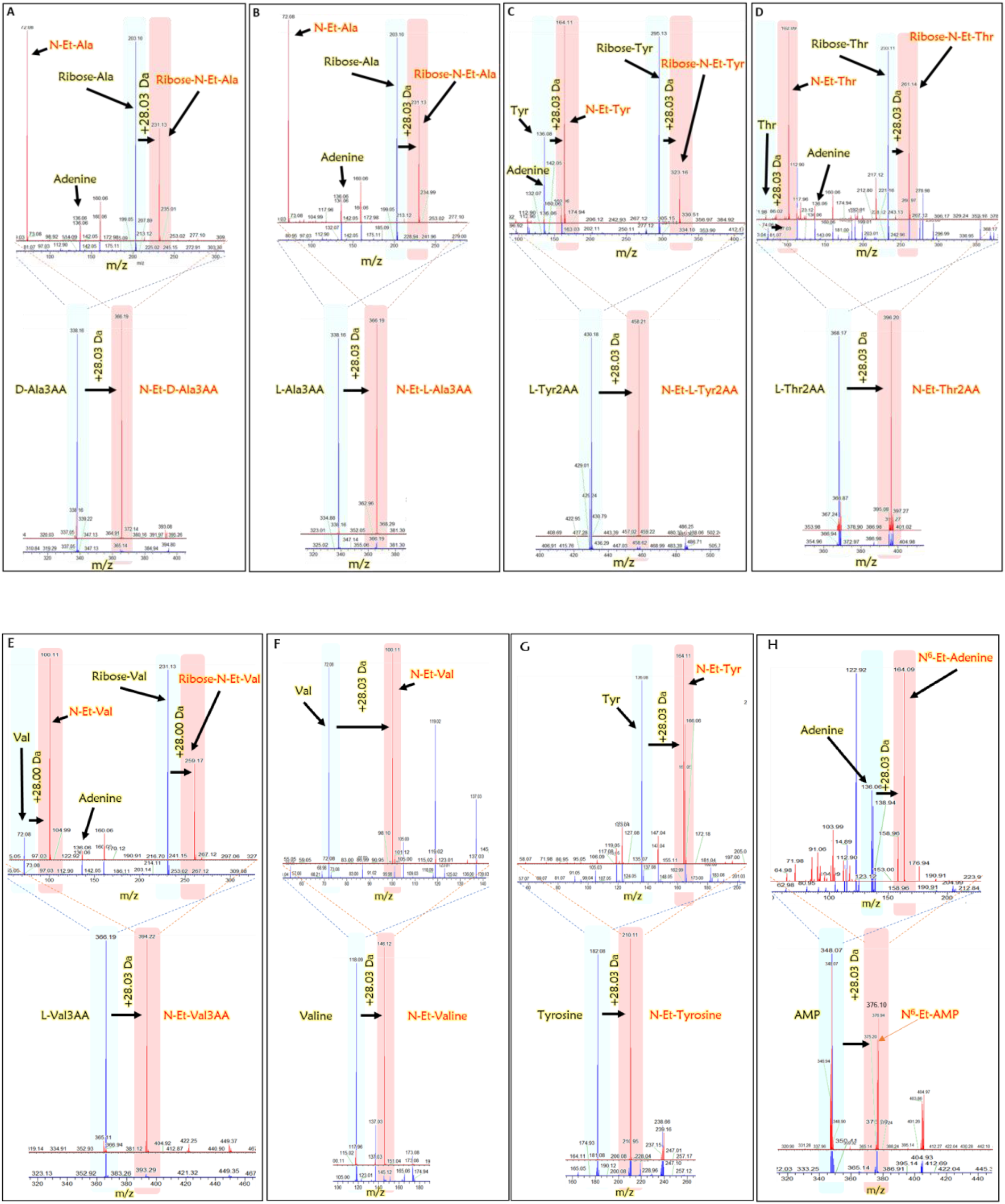
Acetaldehyde modification is independent of aa-tRNA chirality and side-chain. Mass spectrometry analysis showing ethyl modification on substrate analogues of L and D-Ala-tRNA (L and D-Ala3AA), L-Tyr-tRNA (LY2AA), Thr-tRNA (L-Thr2AA) and L-Val-tRNA (L-Val3AA); on amino acids valine and tyrosine and nucleotide AMP (unmodified and modified compound mass spectrum (depicted in blue and red respectively). Fragmentation confirms modification occurs only on the amino acid, not on tRNA of aa-tRNA (fragments depicted in fig. S2 are labelled to their corresponding peaks).

**Extended Data Fig 4:**
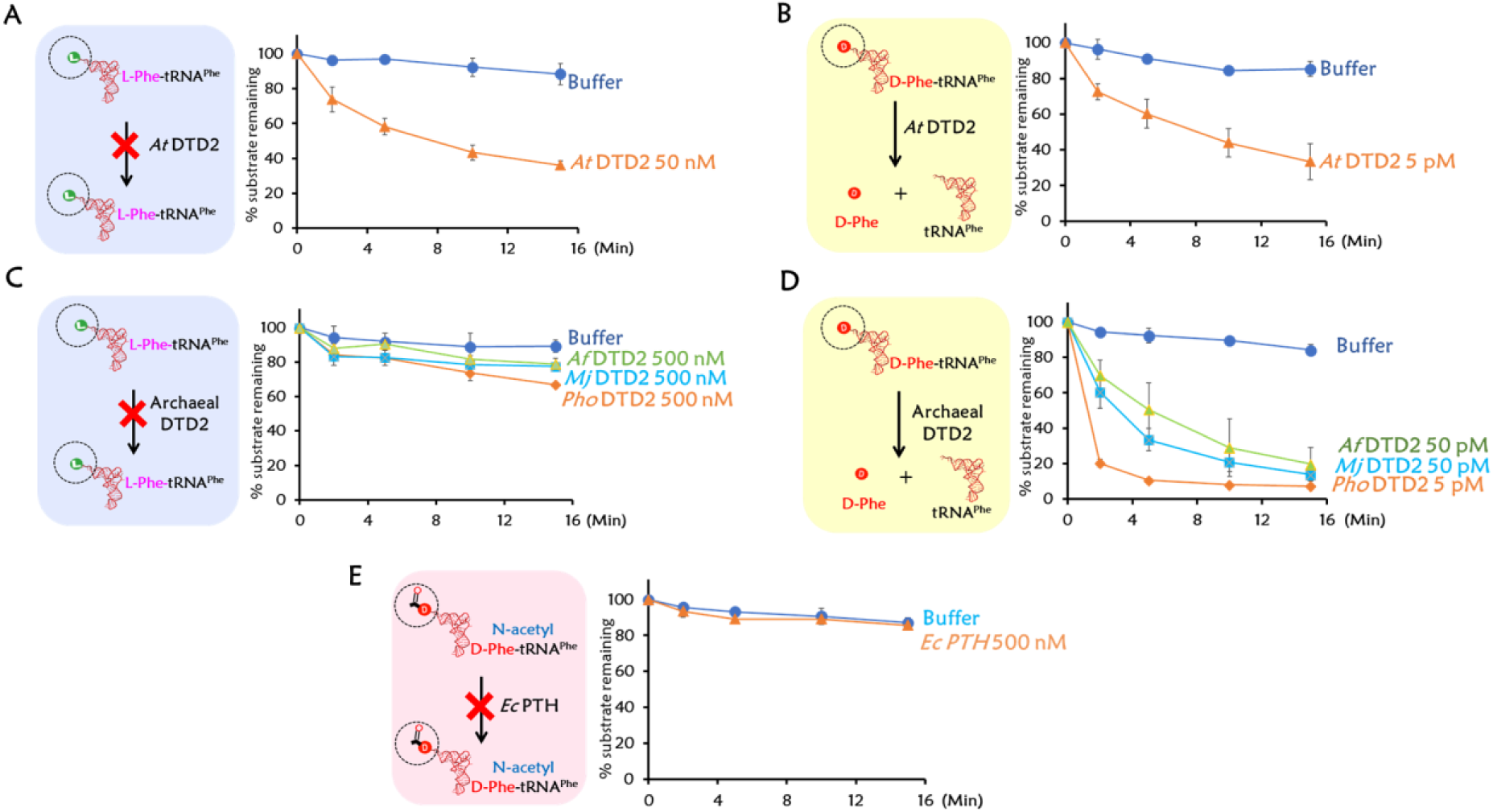
L-Phe-tRNA^Phe^, D-Phe-tRNA^Phe^ and N-acetyl-D-Phe-tRNA^Phe^ deacylations. Deacylations of L-Phe-tRNA^Phe^ (*At*) (A) and D-Phe-tRNA^Phe^ (*At*) (B) with *At* DTD2. Deacylations of L-Phe-tRNA^Phe^ (*Pho*) (C) and D-Phe-tRNA^Phe^ (*Pho*) (D) with archaeal DTD2s (*Pho, Mj and Af*). (E) *Ec* PTH deacylations on N-acetyl-D-Phe-tRNA^Phe^ (*Ec*).

**Extended Data Fig 5:**
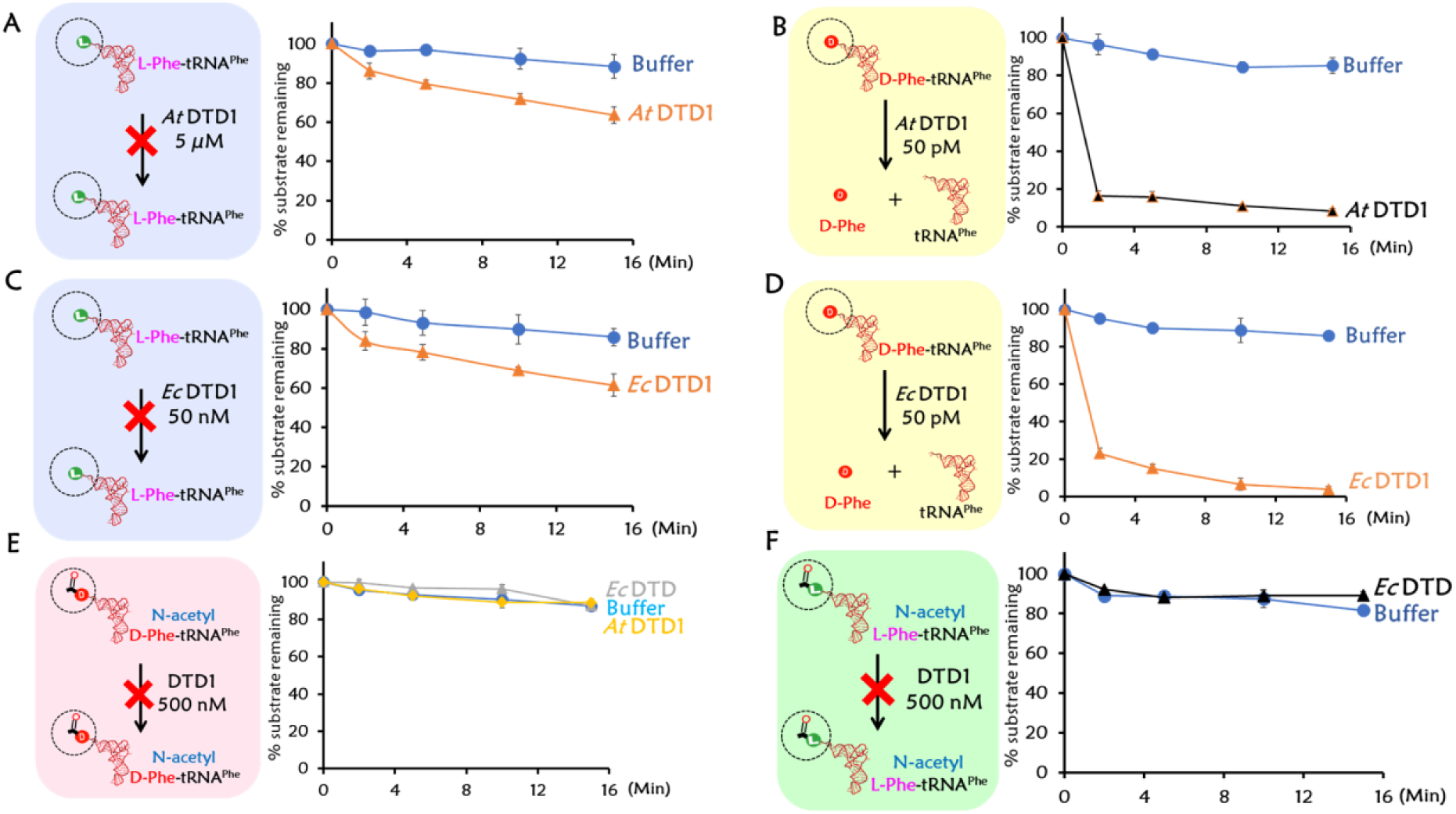
DTD1 deacylations with L- and D-Phe-tRNA^Phe^ and N-acetyl-L- and D-Phe-tRNA^Phe^. Deacylations of L-Phe-tRNA^Phe^ (*At*) (A), D-Phe-tRNA^Phe^ (*At*) (B) with *At* DTD1. *Ec* DTD1 deacylations of L-Phe-tRNA^Phe^ (*Ec*) (C), D-Phe-tRNA^Phe^ (*Ec*) (D) with *Ec* DTD1. N-acetyl-D-Phe-tRNA^Phe^ (*Ec*), (*At*) (E) and N-acetyl-L-Phe-tRNA^Phe^ (*Ec*), (*At*) (F) deacylations with *Ec* and *At* DTD1.

**Extended Data Fig 6:**
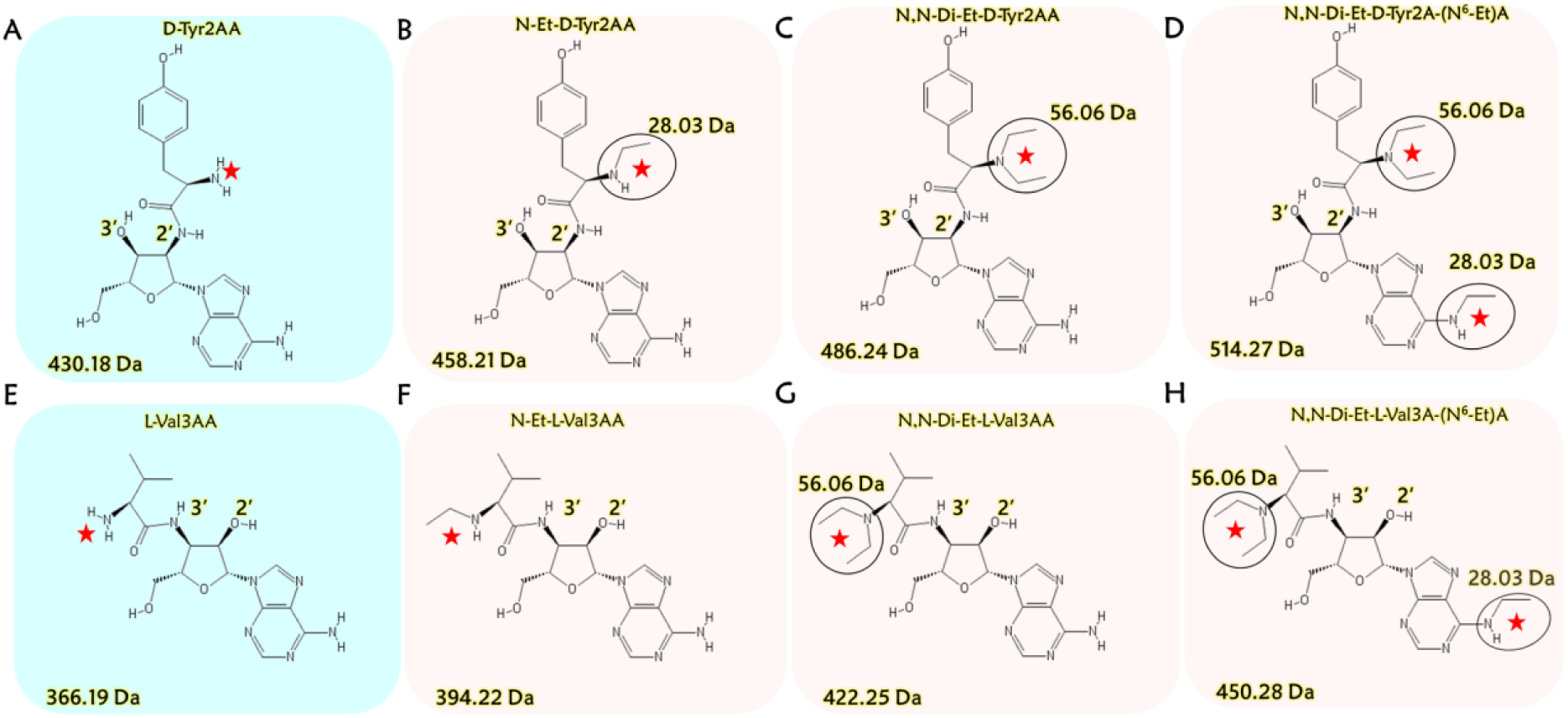
Structures and their expected molecular weights of (A) D-Tyr2AA, (B) N-ethyl-D-Tyr2AA (N-Et-D-Tyr2AA), (C) N, N-diethyl-D-Tyr2AA (N,N-Di-Et-D-Tyr2AA), (D) N, N,N-diethyl-D-Tyr2A(N^6^-ethyl)A (ethyl modification of adenine) (N,N-Di-Et-D-Tyr2A(N^6^-Et)A),, (E) L-Val3AA, (F) N-ethyl-L-Val3AA (N-Et-L-Val3AA), (G) N, N-diethyl-L-Val3AA (N,N-Di-Et-L-Val3AA), and (H) N, N-diethyl-L-Val3A(N^6^-ethyl)A (ethyl modification of adenine) (N,N-Di-Et-L-Val3A(N^6^-Et)A).

**Extended Data Fig 7:**
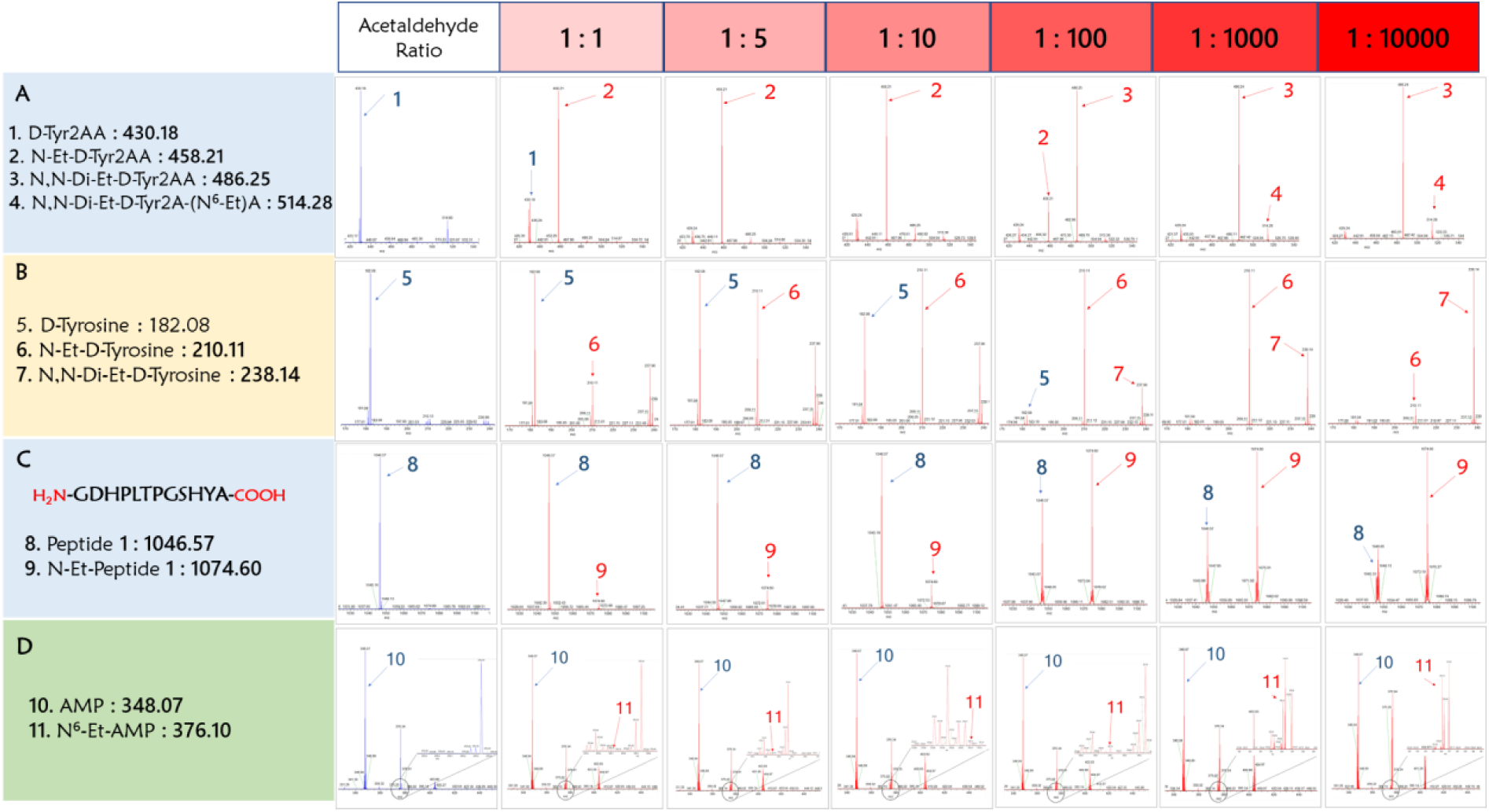
Mass spectrometry data showing hypersensitivity of D-Tyr2AA. (A) Modification on D-Tyr2AA (A), D-Tyrosine (B), peptide 1 (GDHPLTPGSHYA) (C) and AMP (D) at different acetaldehyde concentrations.

**Extended Data Fig 8:**
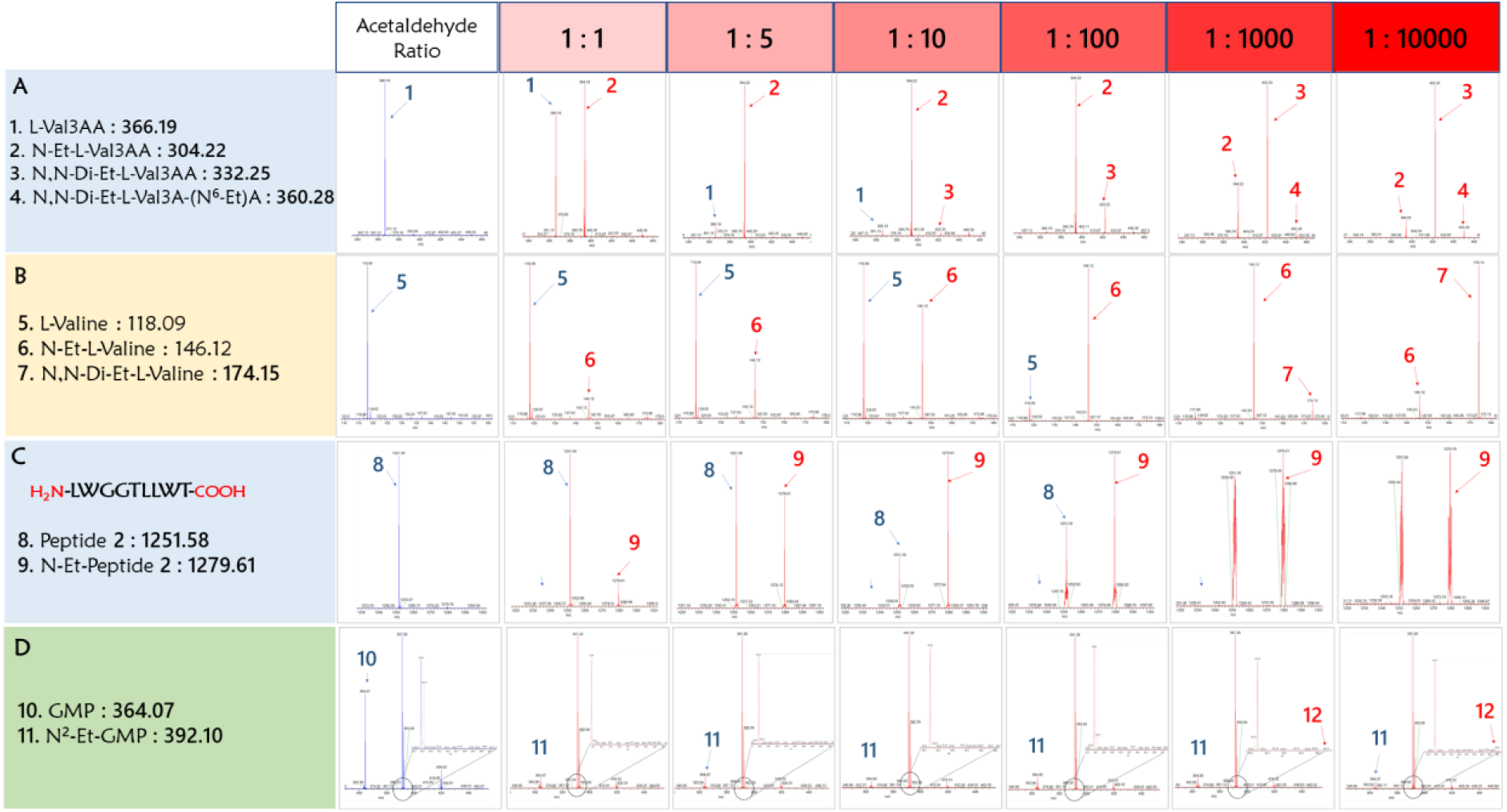
Mass spectrometry data showing hypersensitivity of L-Val3AA. (A) Modification on L-Val3AA (A), L-Valine (B), GMP (C) and peptide 2 (LWGGTLLWT) (D) at different acetaldehyde concentrations.

**Extended Data Fig 9:**
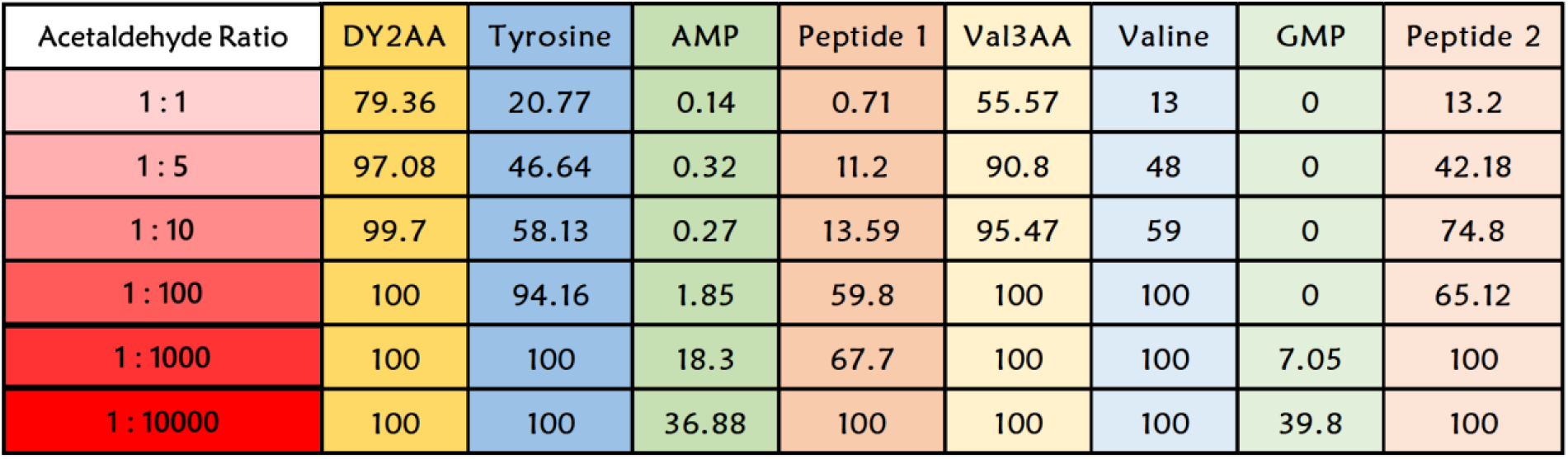
Percentage of modification is calculated from mass spectrometry data represented in figure S4 and S5. Ratios of corresponding peaks observed in mass spectrometry results were used to calculate the percentage modification for respective samples.

**Extended Data Fig 10:**
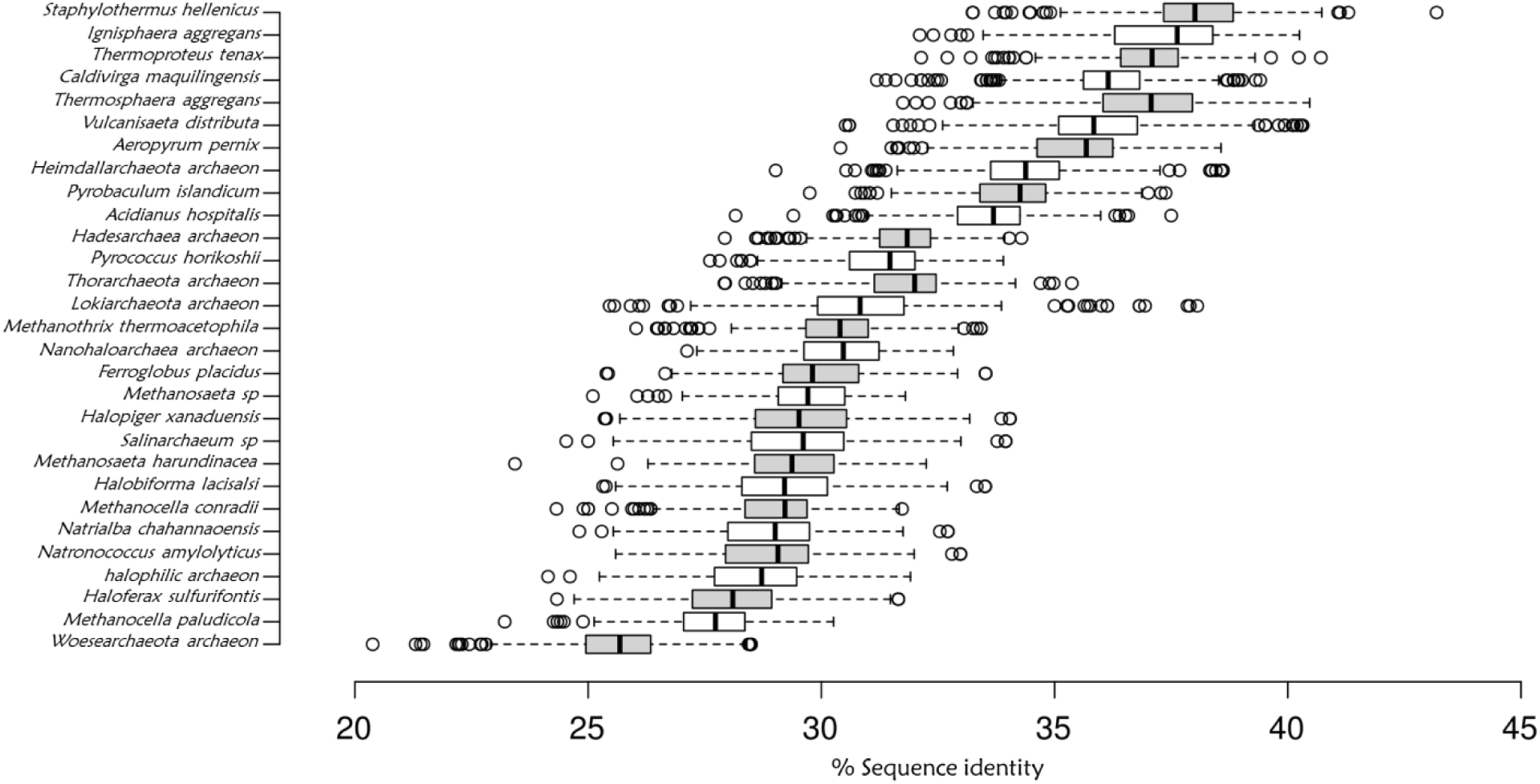
Boxplot displays the percentage of sequence identities of archaeal TopVIB with the plant Top6B. Sequence identity values were obtained from multiple sequence alignment with ClustalO of ∼290 Plant and ∼500 archaeal TopVIB sequences. Only a few representatives were plotted in boxplot (a complete list is provided in XL-sheet). The horizontal axis represents sequence identities with all 290 plants and the vertical axis shows the representative archaeal sequences compared with all the plants.

**Extended Data Fig 11:**
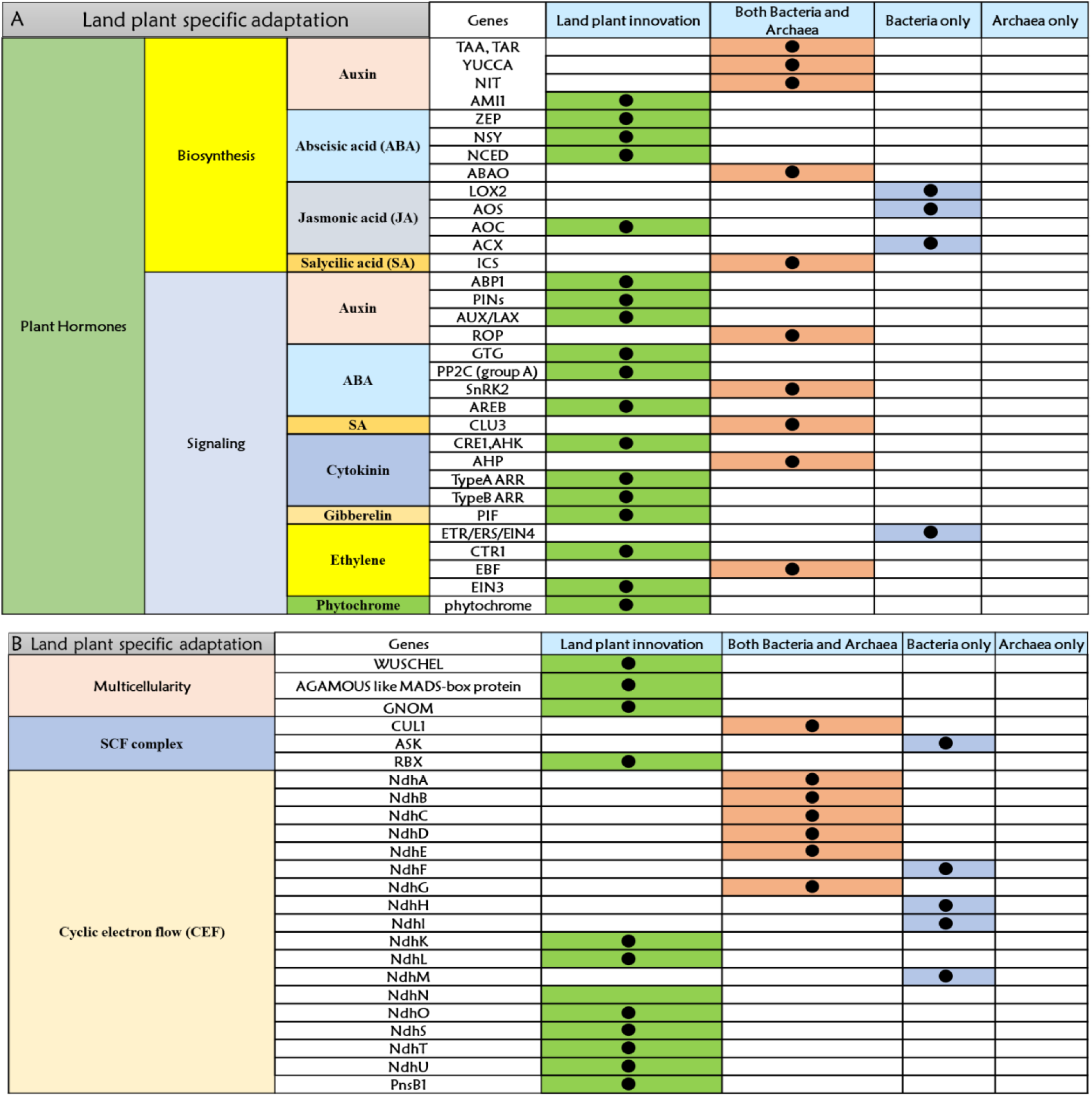
Origins of land plant-specific genes appeared in *Klebsormidium*. Origin of genes responsible for (A) Biosynthesis and signalling of various plant hormones (B) multicellularity, SCF complex and cyclic electron flow in plants.

**Table S1.**
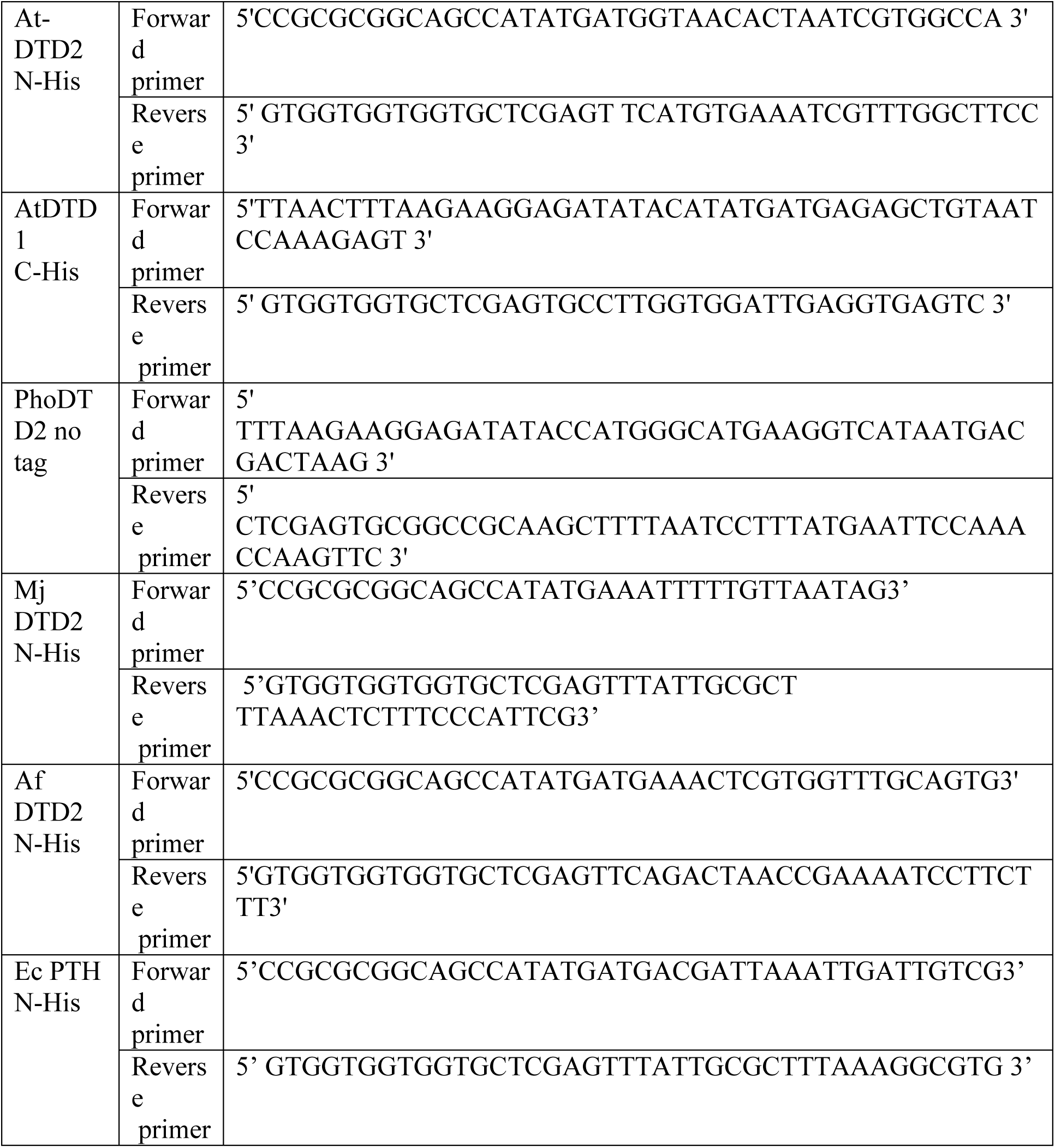
List of primers used in this study.

